# Circadian clock neurons maintain phase control over daily behavioral activity patterns under diverse environmental conditions

**DOI:** 10.1101/2020.06.02.128918

**Authors:** Clara Lorber, Ralf Stanewsky, Angélique Lamaze

## Abstract

Proper timing of rhythmic locomotor behavior is the consequence of integrating environmental conditions and internal time within the circadian clock. The 150 clock neurons in the *Drosophila melanogaster* brain are organized in various clusters, controlling different aspects of the daily activity rhythms. For example, during regular 12 hr light : 12 hr dark cycles at constant temperature (LD), so called Morning (M) neurons control the activity peak in the morning, while Evening (E-) neurons regulate the activity increase at the end of the day. During the remaining times of day and night, flies are inactive, giving rise to the crepuscular behavior observed in LD. Here, we investigate if the same neuronal groups also control behavioral activity under very different environmental conditions of constant light and temperature cycles (LLTC). While the morning activity is completely absent in LLTC, a single pronounced activity peak occurs at the end of the thermophase. We show that the same E-neurons operating in LD, also regulate the evening peak in LLTC. Interestingly, neuronal activity of E-neurons is inversely correlated with behavioral activity, suggesting an inhibitory action on locomotion. Surprisingly, the E-cells responsible for synchronization to temperature cycles belong to the clock neurons containing the circadian photoreceptor Cryptochrome, previously suggested to be more important for synchronization to LD. Our results therefore support a more deterministic function of the different clock neuronal subgroups, independent of specific environmental conditions.

**Significance statement:** Master circadian clocks in the brains of mammals and fruit fly are composed of neurons expressing varying types of neuropeptides and transmitters. This diversity along with anatomical differences indicate diverse functions of different clock neurons. In *Drosophila*, so-called Morning (M) and Evening (E) neurons control locomotor activity at the respective time of day during normal day/night (LD) cycles. Recent reports point to a certain degree of plasticity with regard to circadian clock neuron function, depending on specific environmental conditions. Here we show that one neuronal group, the E-neurons, instead behave as if hard-wired to their output targets. Surprisingly they direct activity to occur during the evening both under LD conditions, as well as during temperature cycles in constant light.

## Introduction

An important function of the circadian clock is to maintain the synchronization of the organism with its ecological niche. To stay on time, the circadian clock is reset everyday by light changes as well as temperature oscillations. These environmental inputs are called ‘Zeitgeber’. Interestingly, while regular temperature cycles are able to synchronize circadian clocks, ambient constant temperature does not influence their free-running period, a phenomenon known as temperature compensation(Pittendrigh 1954; Yoshii et al. 2010). *Drosophila melanogaster*, when placed in a 12h-12h light-dark cycle and constant 25°C (LD), present two peaks of locomotor activity: one starting just a few hours before lights-on and one a few hours before lights-off. These two peaks are controlled by two distinct neuronal oscillators(Grima et al. 2004; Stoleru et al. 2004). However, locomotor activity patterns are highly plastic and responsive to the current environment. For example, at cooler temperature (18°C) the morning peak is not present anymore while the evening peak is advanced compared to milder temperatures(Majercak et al. 1999). On the other hand, at warm temperatures (≥29°C) the evening peak is delayed, while the morning peak is advanced (Majercak et al. 1999). However, because the circadian clock is temperature compensated, these phase shifts of behavioural peaks in response to temperature changes are not due to changes of the free-running period. Hence, the integration of both time information and environmental parameters sets the phase of locomotor activity.

The *Drosophila* brain clock is composed of about 150 clock neurons subdivided by their anatomical position. The circadian blue-sensitive photoreceptor CRYPTOCHROME (CRY) is expressed in about half of the clock neurons(Yoshii et al. 2008). Notably, CRY is expressed in the two neuronal oscillators involved in regulating the locomotor activity pattern in LD. In the absence of a functional visual system, flies are entrained to light due to the presence of CRY and vice versa, while in the absence of both the visual system and CRY the brain clock becomes insensitive to light entrainment(Helfrich-Förster et al. 2001). However, CRY also promotes neuronal activity when activated by light independently of its clock resetting function(Fogle et al. 2011).

Less is known about how the circadian network integrates temperature input for their entrainment and the ambient environment to phase the locomotor output. It has been proposed that the CRY^+^ clock neurons are more sensitive to light while the CRY^-^ neurons are more sensitive to temperature input(Yoshii et al. 2010; Harper et al. 2016). Nevertheless, the behaviour is different between temperature cycles (TC 25°C-16°C) in constant darkness (DDTC) and in constant light (LLTC). In DDTC flies present a peak of activity in the first half of the thermophase while in LLTC they present a peak at the end of the warm phase(Gentile et al. 2013). This suggests that different oscillators control the locomotor behaviour during temperature cycles depending on the light condition but also the range of temperatures used(Gentile et al. 2013). The output of the morning cells is inhibited by light via the visual system (Picot et al. 2007; Murad et al. 2007). In contrast, the output of the evening cells is inhibited by darkness(Grima et al. 2004), suggesting a potential role of the evening cells in controlling the rhythmic behaviour in LLTC.

Here, we show that the evening cells are required for controlling the locomotor behaviour during temperature cycles in constant light, suggesting a deterministic role of this oscillator. However, these CRY^+^ cells are not required for temperature entrainment of the other clock neurons including the CRY^-^ cells suggesting independent temperature entrainment pathways within the circadian network. Finally, we found that the LNd CRY^+^ neurons have their lowest neuronal activity when the flies are the most active, suggesting that the evening cells inhibit locomotion.

## Material and methods

### Fly strains

Flies were raised in a 12 h:12 h light dark (LD) cycle on a medium containing0.7% agar, 1.0% soy flour, 8.0% polenta/maize, 1.8% yeast, 8.0% malt extract, 4.0% molasses, 0.8% propionic acid, and 2.3% nipagin. at 25°C and 60% relative humidity. The following fly lines were used in this study:; *Mai179-GAL4/CyO* (Grima et al. 2004),; *cry[19]-GAL4*(Picot et al. 2007), *spLPN-GAL4* (*R11B03-p65.AD; R65D05-GAL4.DBD*)(Sekiguchi et al. 2020) was provided by Taishi Yoshii,; *DvPdf-GAL4* (Bahn et al. 2009), *R16C05-Gal4*(BL69492) (Schubert et al. 2018), *Clk4.1M-GAL4*(Zhang, Chung, et al. 2010; Zhang et al. 2010), *Clk9M-Gal4*(Kaneko et al. 2012), *Pdf-GAL4*(Renn et al. 1999), *Clk856-GAL4*(Gummadova et al. 2009), *UAS-TNTG*(Sweeney et al. 1995) (BL28838),; *UAS-cyc*^*DN*^ (Tanoue et al. 2004), *Pdf-GAL80* (BL80940), *UAS-CD4TdTom* (BL35837), *UAS-CaMPARI*^*V398D*^ (Fosque et al. 2015) (BL58762). *Mai179-GAL4, UAS-TNTG, UAS-cyc*^*DN*^, *Clk9M-GAL4*, and *Clk4.1-Gal4* have been outcrossed for 5 generations to *iso31* to standardize the genomic background (Liu et al. 2014).

### Behavior

Analysis of locomotor activity of male flies was performed using the Drosophila Activity Monitoring System (DAM2, Trikinetics Inc., Waltham, MA, USA) with individual flies in recording tubes containing food (2% agar, 4% sucrose). Briefly, DAM2 activity monitors containing LD-entrained flies were placed inside a light- and temperature-controlled incubator (Percival Scientific Inc., Perry, IA, USA). Fly activity was monitored for at least 11 days with the first 2 days in LD25°C (700-1000 lux generated by 17W F17T8/TL841 cool white Hg compact fluorescent lamps, Philips) followed by three days in LL 25°C and then LLTC 25°C-16°C with a shift delayed by 5h relative to the previous LD. The LD behavior analysis was performed on the last day of LD, while LLTC behavior was analyzed on the 6^th^ day of LLTC.

Plotting of actograms was performed using a signal-processing toolbox(Levine et al. 2002) implemented in MATLAB (MathWorks, Natick, MA, USA). The 24h locomotor activity plots were performed using a custom excel macro (Lamaze et al. 2017). Activity was averaged into 30 minute bins and normalized to the maximum individual activity level. The median of this normalized activity was plotted, because it is a more representative parameter of behavioral synchrony compared to the mean. To measure the slope in LD, we manually determined the latest time point of the minimum median (t_min_) and we calculated for each fly the derivative of the line between the time point 691 (the last half hour before light-off) and t_min_ : Slope=(Act_691_-Act_min_)/(t_691_-t_min_). In LLTC, because the evening peak is advanced and therefore we can see the maximum activity, we instead determined the time point of the maximum median level and measure the slope as follow: Slope=(Act_max_-Act_min_)/(t_max_-t_min_). The box plots were made using Excel. The statistical tests were performed using the freely available Estimation Statistics BETA(Ho et al. 2019).

### Immunostaining

Adult male *Drosophila* brains were immuno-stained as described previously [Liu et al, Neuron 2014]. Briefly, brains were fixed in 4% paraformaldehyde for 20 min at RT, and blocked in 5% goat serum for 1 h at RT. Primary antibodies used were as follows: rabbit anti-DsRed (Clontech) – 1:2000; mouse anti-PDF (Developmental Studies Hybridoma Bank, DSHB) – 1:2000; mouse anti-Bruchpilot (nc82, DSHB) – 1:200; chicken anti-GFP (Invitrogen) – 1:1000 – rabbit anti-PER(Stanewsky et al. 1997) 1:15000. Alexa-fluor secondary antibodies (goat anti-rabbit 555, goat anti-chicken 488, goat anti-mouse 488; Invitrogen) were used at 1:2000 except for labeling anti-BRP where goat anti-mouse 647 where a dilution of 1:500 was used. Confocal images were taken using an inverted Leica LSP8. PER signal intensity was quantified using an 40x oil-objective and imageJ. Pixel intensities were normalized to the background after background subtraction [(signal-background)/background] using Excel.

### Calcium imaging

CaMPARI experiments were performed as described previously (Lamaze et al. 2018). Briefly, 1-3 days old flies were entrained in LLTC. Several days prior to the experiment, flies were transferred to individual glass tubes to facilitate isolation without CO_2_ or cold-induced anesthesia. On the experimental day, brains were dissected in HL3.1 buffer (70 mM NaCl, 5 mM KCl, 20 mM MgCl2, 1.5 mM CaCl2, 10 mM NaHCO3, 5 mM Trehalose, 115 mM sucrose, 5 mM HEPES, pH 7.2). To induce photo-conversion, brains were exposed to UV using the high power mercury lamp from the microscope for 1 min. The less calcium-sensitive V398D variant of CaMPARI was used to increase the dynamic range of photo-conversion (Fosque et al. 2015).

## Results

### The evening cells control the evening activity during temperature cycles in constant light

The presence of only one clock-controlled locomotor activity peak in LLTC suggests that only one oscillator output is active under this condition. Since the phase of this activity peak occurs in the second half of the thermophase, we hypothesized that the evening cells regulating behavior in LD might also play a role in regulating behavior during temperature cycles.

To test this hypothesis, we first stopped the neuronal activity of the morning and evening cells using *Mai179*-gal4(Grima et al. 2004) and UAS-TNTG (Sweeney et al. 1995). *Mai179* is expressed in the morning cells (sLNv) and the evening cells (5^th^ LNv, 3 LNd CRY^+^ and 2 DN1a) as well as many other non-clock peptidergic neurons(Grima et al. 2004; Picot et al. 2007). Stopping the neuronal activity and at the same time rescuing the molecular clock in these cells, strongly decreases the amplitude of the evening anticipation in LD without affecting the molecular clock(Johard et al. 2009). Consistent with this observation, *mai179 > TNTG* flies have a strongly dampened evening peak confirming both, that the UAS-TNTG tool is working and that the *Mai179* cells drive evening anticipatory behavior in LD (Fig. 1A, B). Interestingly, we observed an even stronger phenotype in LLTC (Fig. 1A) with a complete absence of evening anticipation (Fig. 1C). As a measure for the synchrony among flies and the amplitude of the evening peak, we quantified the behavioral activity increase in the 2^nd^ half of the thermophase before the temperature drop at ZT12. The extend of this anticipatory behaviour was defined as the ‘slope’ (see Material and Methods for details). Briefly, if the median of the slope is not significantly different from 0, flies are considered incapable of showing any synchronized anticipatory behavior. Controls show a slope ≥0.4 in LD and around 0.3 in LLTC. In LD, Mai179 > TNTG have a slope significantly different from 0 but also significantly lower than controls (Fig. 1D), suggesting that, although the amplitude is dampened, they are still synchronized. In contrast, in LLTC, the median of the evening slope was not significantly different from 0 (Fig. 1E), indicating that the flies do not anticipate the temperature transition. To confirm the role of the CRY^+^ morning and evening cells, we used the *cry[19]*-Gal4 driver which is expressed in the same clock cells as *Mai179*, but absent from the other non-clock peptidergic neurons(Picot et al. 2007). *cry[19] > TNTG* flies showed similar behavior as *Mai179 > TNTG* flies, both in LD and in LLTC (Fig. 2). Hence, we conclude that the Mai179^+^ clock neurons mediate temperature synchronization in LL, ruling out a function for the non-clock peptidergic neurons, expressing this driver.

**Figure 1:**
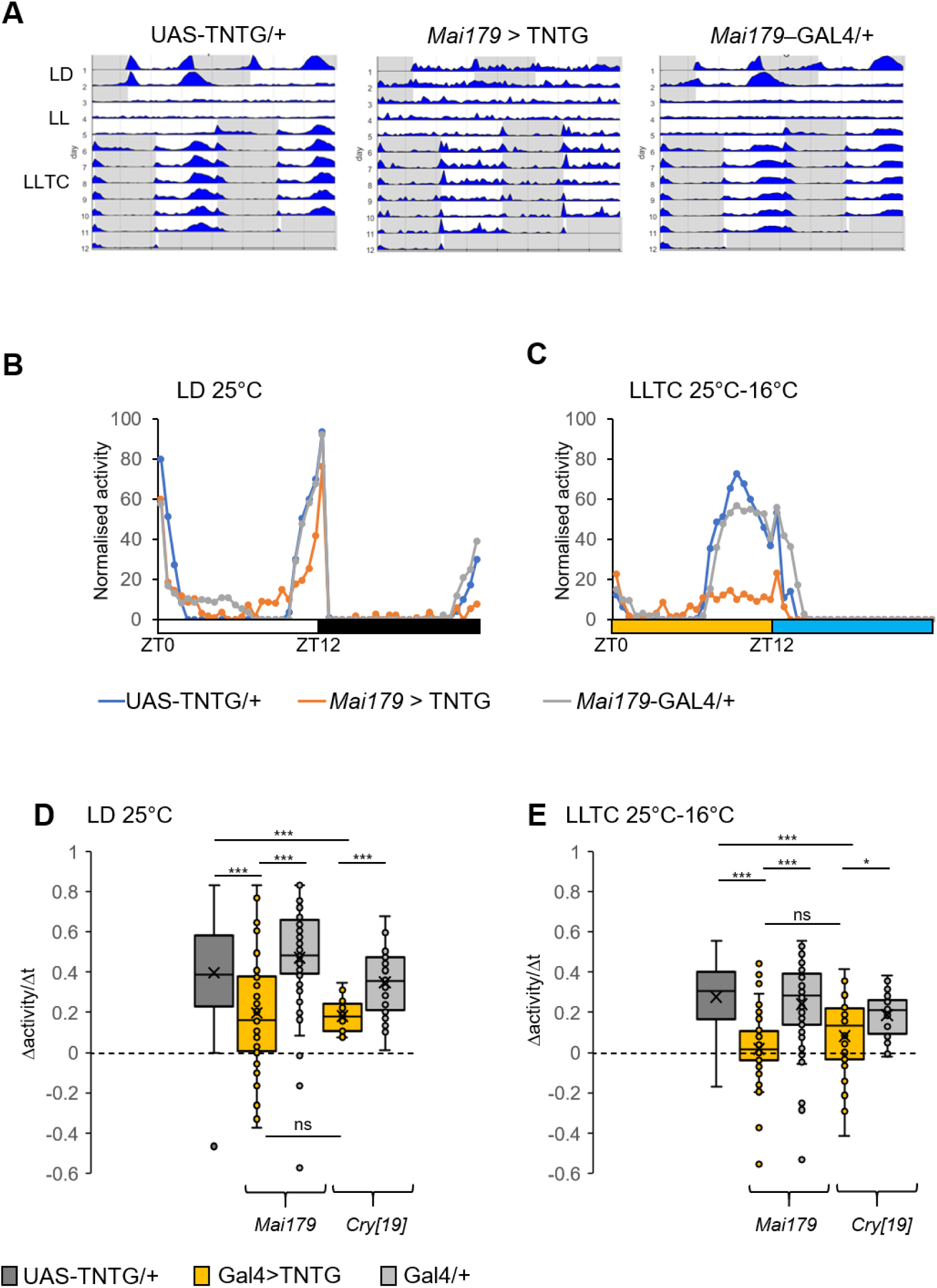
Inhibition of the morning and evening cells stops the rhythm in LLTC. Male flies were loaded in LD 25°C. After 2 days, we switched the environment to LL 25°C for three days then we started the temperature entrainment in LL (LLTC 25°C-16°C) with a 5h delay with the previous LD cycle. A) Group actograms of a representative experiment. N(UAS-TNTG/+): 20, N(*Mai179* > TNTG): 13, N(*Mai179*-GAL4/+): 20. B-C) Median of the normalised locomotor activity during the last day of LD (B) and the 6th cycle of LLTC (C). White bar represents the 12h of light-on, black bar, yellow and blue bars schematise darkness, thermophase and cryophase periods respectively. N(UAS-TNTG/+): 60, N(*Mai179* > TNTG): 54, N(*Mai179*-GAL4/+): 57. D-E) Box plot of the slope of the evening peak in the last day of LD (D) and the 6^th^ cycle of LLTC (E). The slope is measured as followed. In LD, we subtracted the level of activity at the last time point before light-off with the last minimum median divided by the time length. In LLTC we subtracted the level of activity at the first time point of the maximum median activity with the last minimum median divided by the time length. For *Mai179* > TNTG We used the same time points as the controls. N(UAS-TNTG/+): 80, N(*Mai179* > TNTG): 54, N(*Mai179*-GAL4/+): 57, N(*cry[19]* > TNTG): 31, N(*cry[19]*-GAL4/+): 29.Statistical test: Kruskal wallis(Ho et al. 2019). *: p<0.05, **:p<0.005, ***:p<0.001

**Figure 2:**
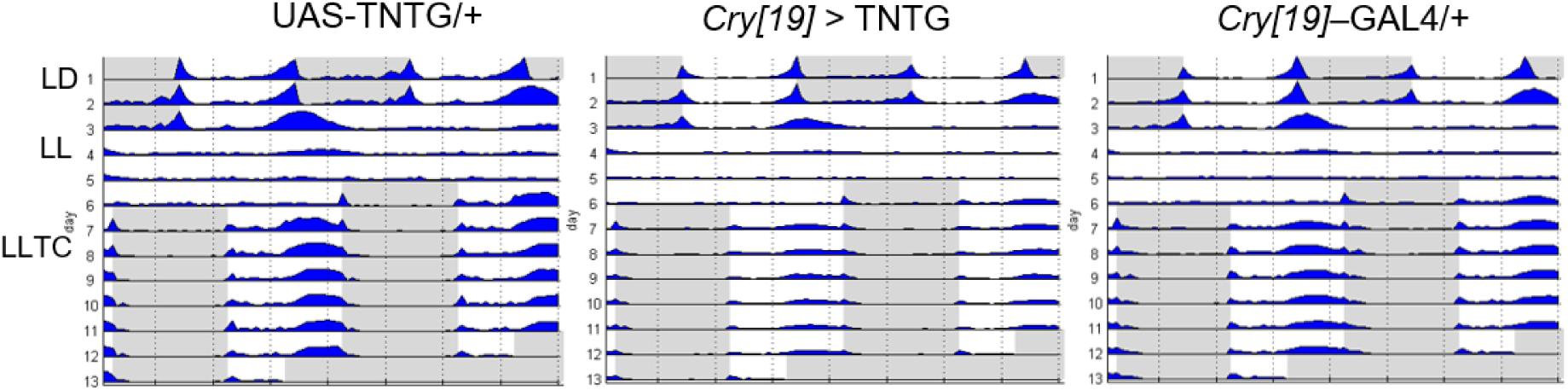
Average actograms. N(UAS-TNTG/+): 18, N(*cry[19]* > TNTG): 32, N(*cry[19]*-GAL4/+): 28

To investigate the potential role of other clock neurons and to distinguish between the role of morning and evening cells, we performed a screen using the most restricted drivers available (Table 1). Decreasing the neuronal activity of non E- or M-clock neurons using TNTG did not affect the evening peak in LLTC (Fig. 3A). Notably, inhibition of candidate neurons important for temperature entrainment such as the DN2 (*Clk9M*-GAL4), DN1p (*Clk4.1M*-GAL4) or the LPN (*spLPN*-GAL4)(Sekiguchi et al. 2020) does not disturb synchronized locomotor activity in LLTC (Fig. 3A). Notably, *Clk9M*-gal4 is expressed in the DN2 as well as in the morning cells (sLNv)(Kaneko et al. 2012). Therefore, the absence of phenotype observed with this driver suggests that the morning cells have a small or no influence on the evening peak in LLTC. Interestingly, *R16C05*>TNTG shows a slight but significant decrease of the evening peak amplitude (Fig. 3A, Fig. 4A). *R16C05*-GAL4 is expressed in the 2 DN1a as well as 2 LNd CRY^+^ neurons(Schubert et al. 2018) (Fig. 4B-C). To confirm that the M cells are not required for synchronization in LLTC, we applied *DvPdf*-gal4, which is expressed in the PDF^+^ (Pigment Dispersing factor) lateral neurons as well as the 3 CRY^-^ LNd, the 5^th^ sLNv, and 1 CRY^+^ and ITP^+^ LNd(Schubert et al. 2018). Expression of the TNTG in these cells does not affect the evening activity in LLTC (Fig. 3B,C) confirming that neuronal activity of M-cells does not play a role in the synchronized locomotor activity in LLTC. Furthermore, inhibition of the 5^th^ and one LNd CRY^+^ (DvPdf-gal4) is not enough to prevent synchronization of locomotor behavior in LLTC. Hence, we can conclude that the strong phenotypes observed with *Mai179-gal4* and *cry[19]-gal4* are caused by blocking the E- and not the M-neurons addressed by these drivers. Nevertheless, we cannot rule out an influence from the morning cells to the evening cells and a minor role for the DN1a.

**Table 1:**
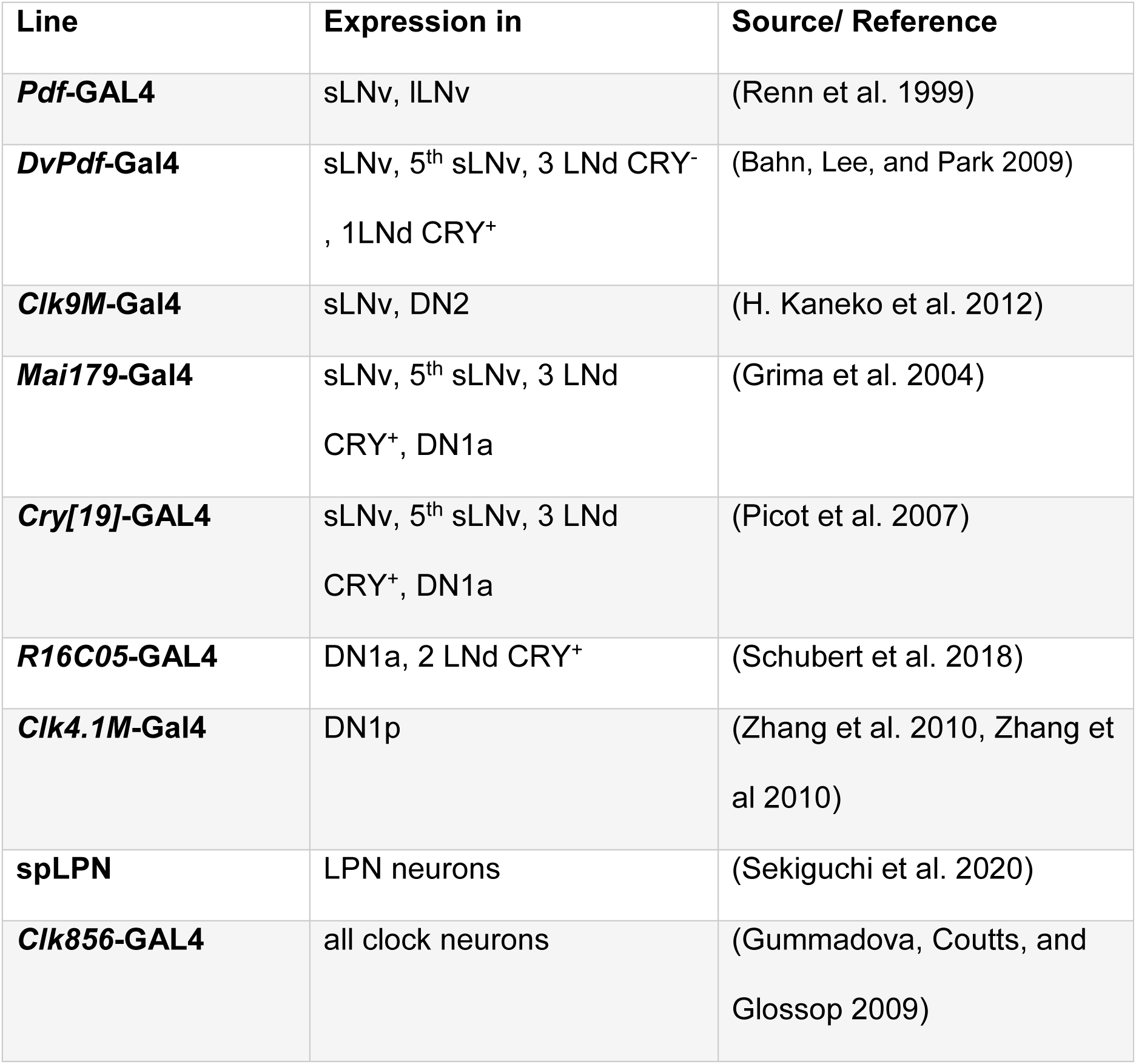
List of *Gal4* drivers and their expression pattern used in this study.

**Figure 3:**
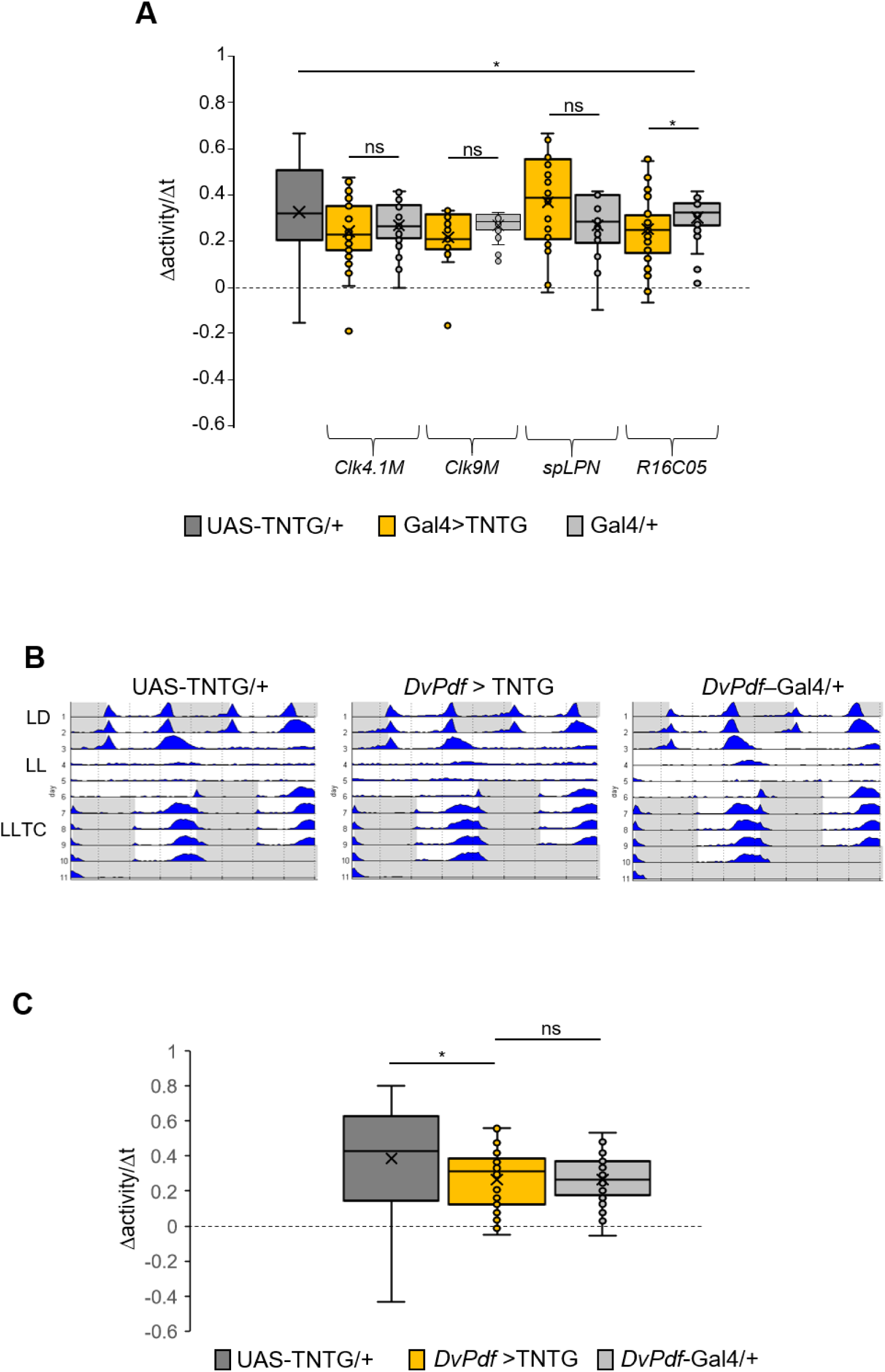
Inhibition of the non-evening cells does not stop the rhythm in LLTC. A) Box plot of the slope of the evening peak in the 6^th^ LLTC cycle. 18≤N≤58. B) Group actograms of a representative experiment. N(UAS-TNTG/+): 20, N(*DvPdf* > TNTG): 20, N(*DvPdf*-GAL4/+): 20. C) Slope of the evening peak during the 6^th^ cycle of LLTC. N(UAS-TNTG/+): 42, N(*DvPdf* >TNTG): 30, N(*DvPdf*-GAL4/+): 40. Statistical test: Kruskal wallis. *: p<0.05, **:p<0.005, ***:p<0.001.

**Figure 4:**
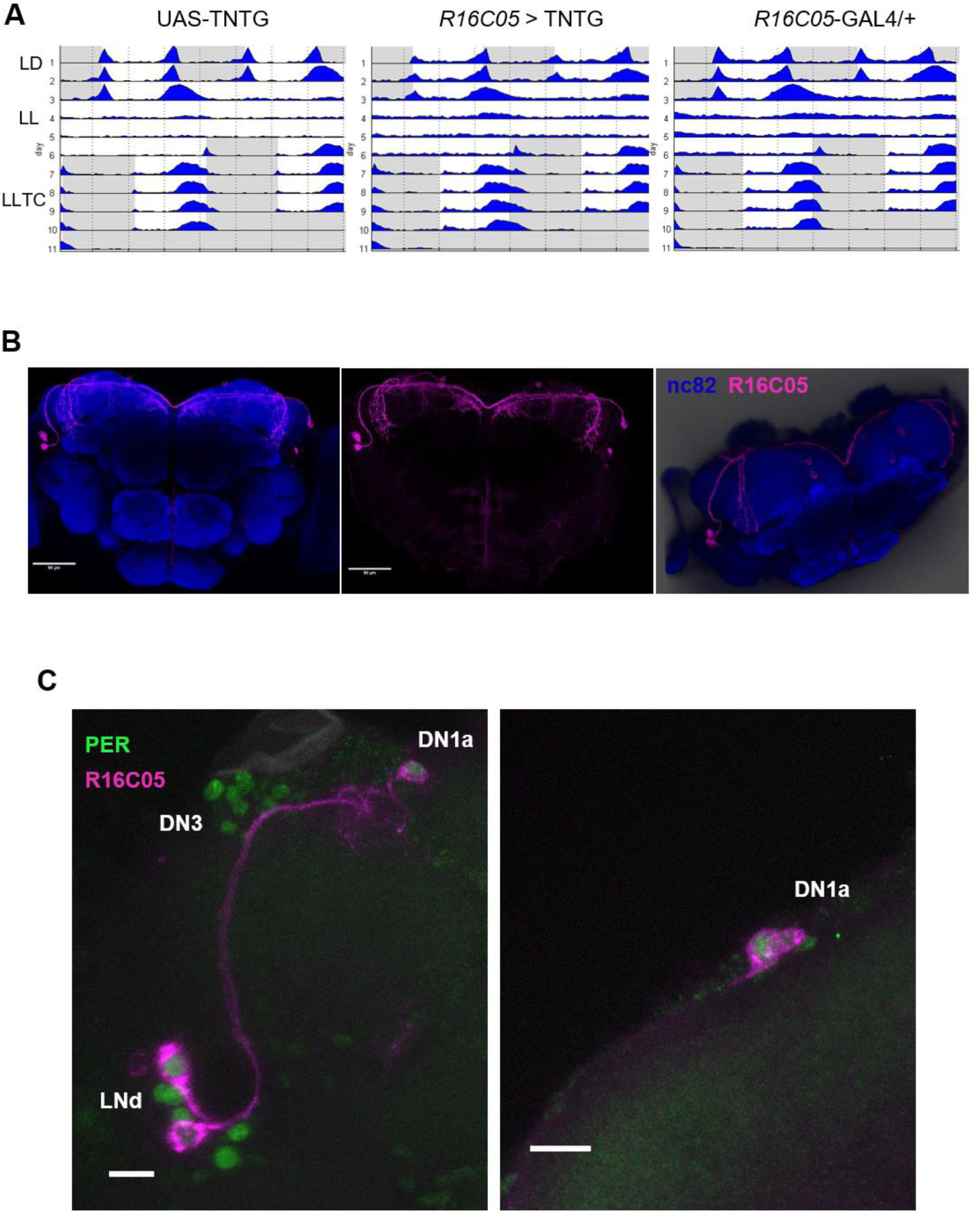
A) Average actograms. B) Projection pattern of R16C05 cells in the fly brain. Genotype: *R16C05* > CD4TdTomato. The neuropiles were labelled with nc82 (blue). The right panel is 3D superior view of the same brain. The DN1a are proximal to the tip of the α,α’ arms of the mushroom bodies. Scale bar: 50µm. C) PER immunostaining (green) in *R16C05* > CD4TdTomato flies. Left picture: dorso-lateral region of the brain. We can see the LNd, the DN3 and the DN1a. Right panel: DN1a. Scale bar: 10µm.

### A functional clock in the E cells is necessary for properly timed evening activity during temperature cycles in constant light

Neuronal activity of the CRY^+^ evening cells is required for a proper locomotor activity pattern in LLTC. This observation is surprising since it was generally assumed that the CRY^-^ clock neurons are relevant for temperature entrainment (Yoshii et al. 2010; Harper et al. 2016; Miyasako et al. 2007). Therefore, we wanted to test if these neurons also require a functional molecular clock to drive locomotor behavior in LLTC.

First, we stopped the clock in the Mai179 CRY^+^ cells (Figure 5A-E). For this, we used the dominant negative UAS-cyc^DN^ genetic tool, previously shown to disrupt rhythms of clock gene expression and behavior (Tanoue et al. 2004; Bulthuis et al. 2019). To confirm the effect of this tool on the molecular clock, we expressed *cyc*^*DN*^ in all clock neurons using *Clk856*-GAL4 (Gummadova et al 2009). During LLTC, we observed a complete absence of synchronized activity with a slope not significantly different from 0 (Fig. 6B), confirming the efficacy of the *cyc*^*DN*^ tool. Interestingly, in *Mai179* > *cyc*^*DN*^ during LD, the evening peak is strongly dampened, but the morning peak is present and comparable to the controls (Fig. 5B-D). This is consistent with previous findings showing that a functional clock in the morning cells, although sufficient, is not required for morning anticipation(Grima et al. 2004; Stoleru et al. 2004; Zhang, Liu, et al. 2010).

**Figure 5:**
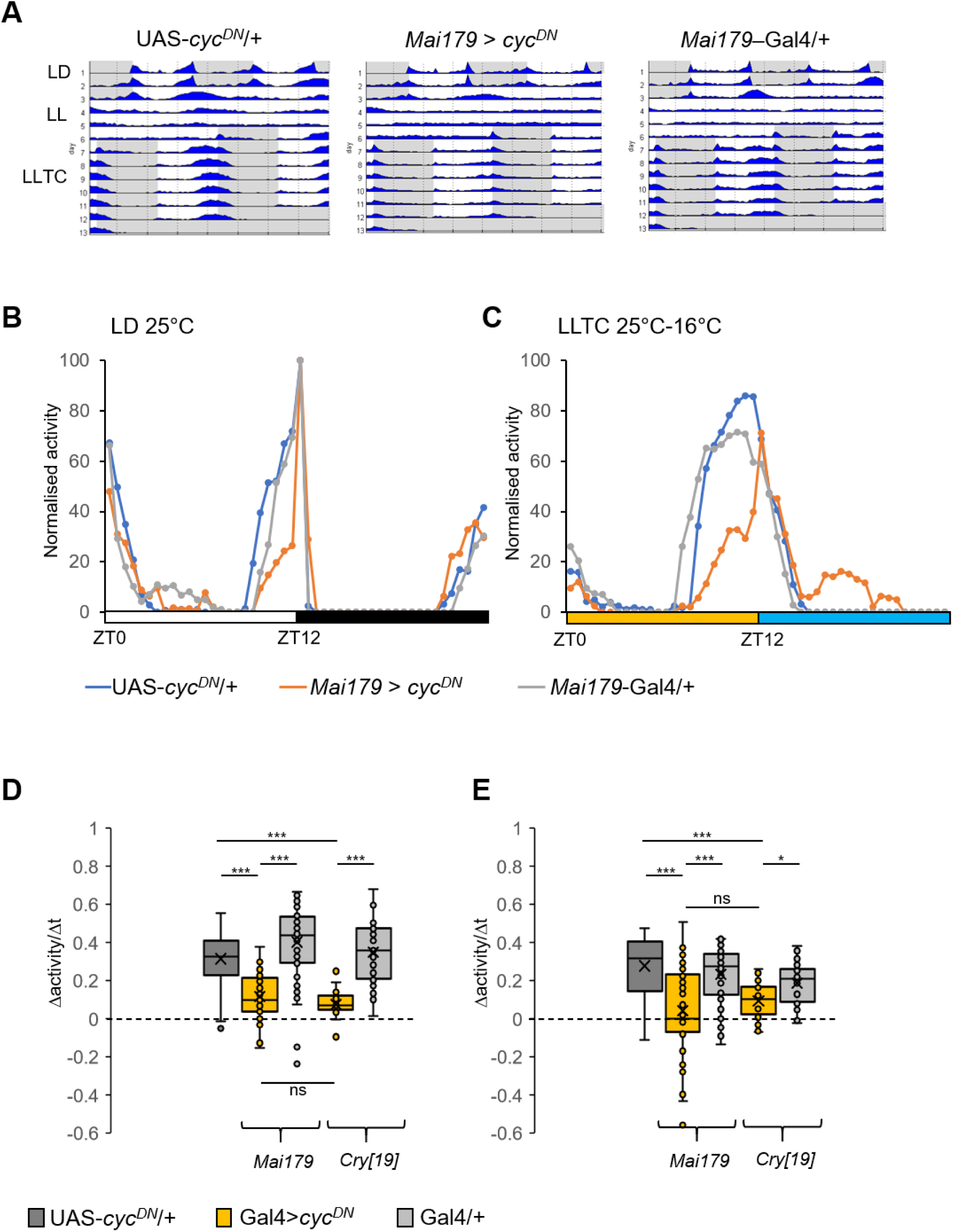
Stopping the clock in the morning and evening cells stops the rhythm in LLTC. A) Group actograms of a representative experiment. N(UAS-*cyc*^*DN*^/+): 19, N(*Mai179* > *cyc*^*DN*^): 21, N(*Mai179*-GAL4/+): 20. B-C) Median of the normalised locomotor activity during the last day of LD (B) and the 6th cycle of LLTC (C). White bar represents the 12h of light-on, black bar, yellow and blue bars schematise darkness, thermophase and cryophase periods respectively. N(UAS-*cyc*^*DN*^/+): 59, N(*Mai179* > *cyc*^*DN*^): 55, N(*Mai179*-GAL4/+): 59. D-E) Box plot of the slope of the evening peak in the last day of LD (D) and the 6^th^ cycle of LLTC (E). N(UAS-*cyc*^*DN*^/+): 75, N(*Mai179* > *cyc*^*DN*^): 55, N(*Mai179*-GAL4/+): 59, N(*cry[19]* > *cyc*^*DN*^): 22, N(*cry[19]*-GAL4/+): 29.Statistical test: Kruskal wallis. *: p<0.05, **:p<0.005, ***:p<0.001.

**Figure 6:**
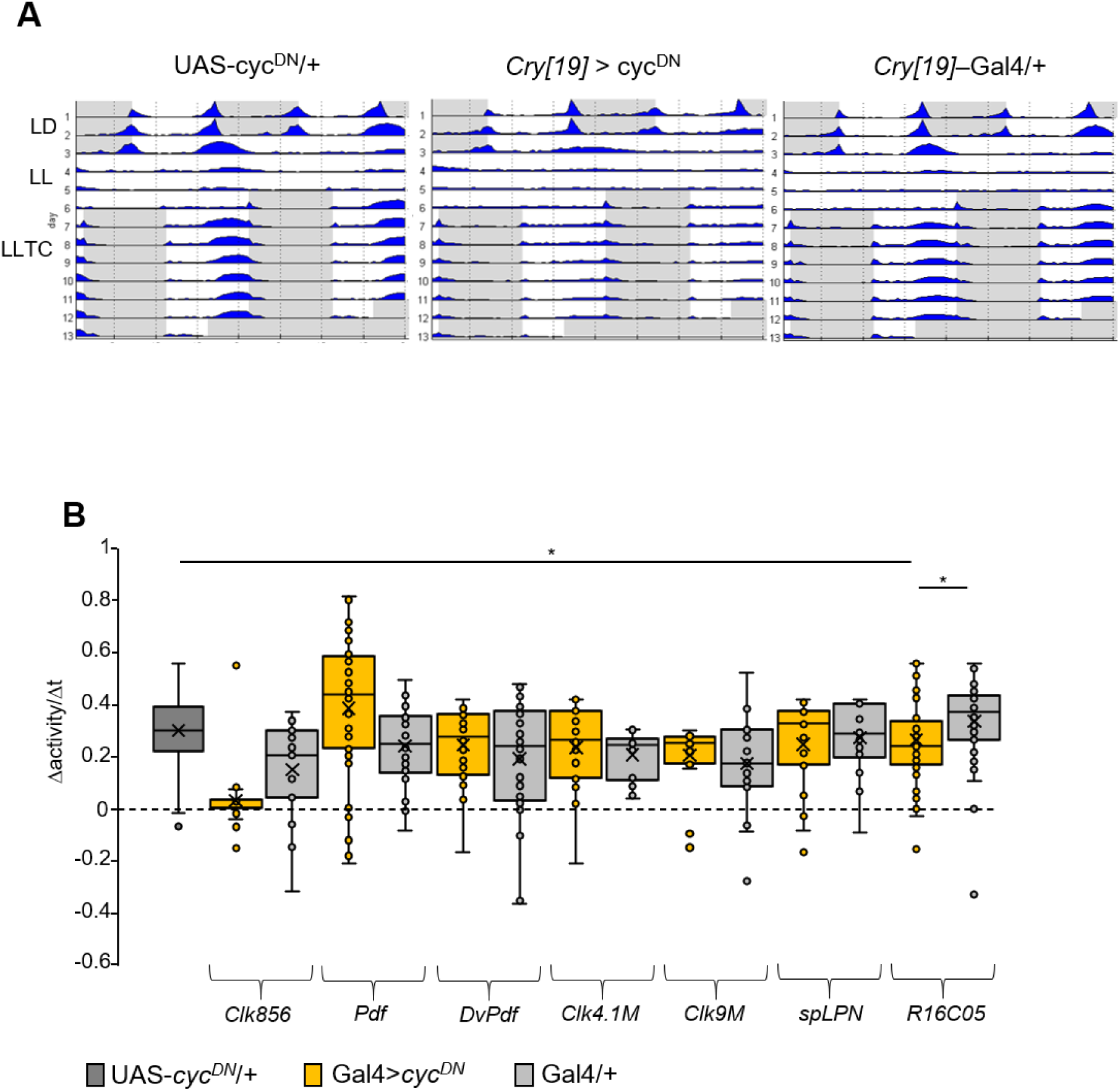
A) Average actograms. N(*UAS-cyc*^*DN*^/+): 16, N(*cry[19]* > *cyc*^*DN*^): 22, N(*cry[19]*-GAL4/+): 28. B) Slope of the evening peak during the 6^th^ cycle of LLTC. N(*UAS-cyc*^*DN*^/+): 153, for the other genotypes 14≤N≤52.

Strikingly, in LLTC, the *Mai179 > cyc*^*DN*^ flies show severely reduced anticipatory behaviour (Fig. 5A,C,E). Although the median of the normalised activity seems to only show a dampening of the evening peak (Fig. 3C), the median of the slope was not significantly different from 0 (Fig. 3E). Similar results were obtained with the *cry[19]*-Gal4 driver (Fig. 5D-E, Fig. 6A), confirming the results obtained with *Mai179*-Gal4. To test for a potential role of non M- and E-cells, we again screened the other clock neuronal groups in LLTC using our set of GAL4 drivers. Consistent with the neuronal silencing results, interfering with clock function in candidate temperature entrainment neurons did not result in a significant reduction of the amplitude of the evening peak (Fig. 6B). Importantly, stopping the clock with any of the drivers expressed in the M-cells did not affect evening anticipation in LLTC (Fig. 6B). Finally, and similar to neuronal silencing, we did observe a slight but significant decrease of the evening peak when we stopped the clock in the DN1a and 2 LNd CRY^+^ using *R16C05*-GAL4 (Fig. 6B).

Although we have shown that a clock in the M-cells is not required for the evening peak in LLTC, we wanted to rule out a potential influence of the morning clock on the evening one. For this, we recombined UAS-*cyc*^*DN*^ with *pdf*-Gal80 and combined these constructs with *Mai179*-GAL4 and UAS-gfp. These flies are expected to have a functional clock in the PDF M-cells, but not in the E-cells. To confirm this, we dissected these flies in LD at ZT2 and determined GFP, PER, and PDF expression. Indeed, PER levels were strongly decreased in the evening cells while the sLNv were PER positive and GFP negative (Fig. 7A). We then measured the locomotor activity of flies expressing *cyc*^*DN*^ in the *Mai179* PDF^-^ cells. We observed a strong dampening of the evening peak as well as a dampening of the slope in both LD and LLTC (Fig. 7B-E). The behaviour in LLTC was essentially indistinguishable from that of *Mai179* > *cyc*^*DN*^ flies without *Pdf-gal80* (Fig. 5C, E). Hence, we conclude that a clock in the sLNv is not sufficient and has no influence on the other Mai179 cells, for the control of the rhythmic locomotor activity in LLTC.

**Figure 7:**
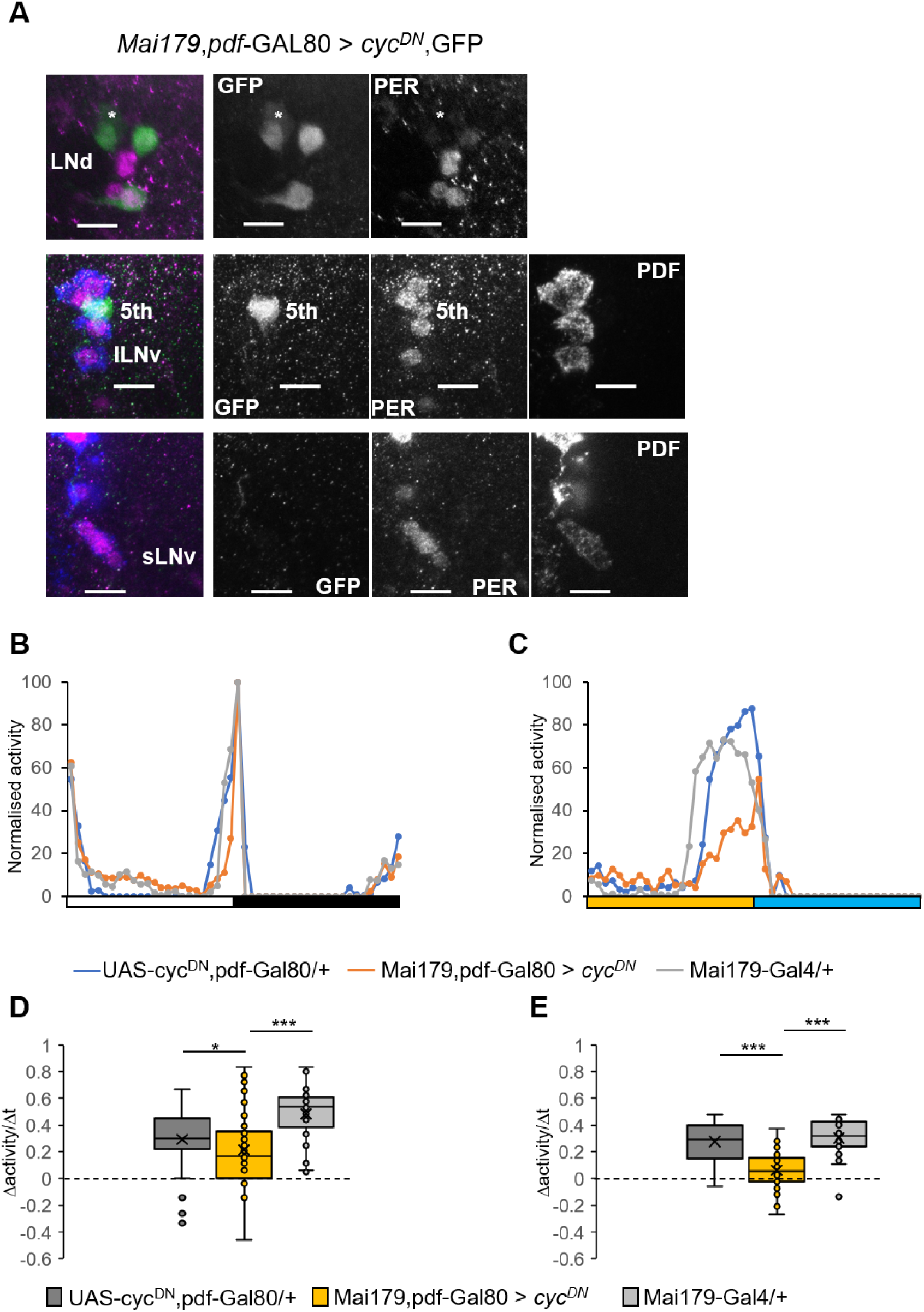
Stopping the clock in the evening cells is sufficient to disturb the rhythm in LLTC. A) Immunostaining of morning and evening cells in LD ZT2. Left panels are the merge images. Green: GFP, magenta: PER, blue: PDF. *: non-clock neuron Mai179^+^ in the LNd region. Scale bar:10µm. B-C) Median of the normalised locomotor activity during the last day of LD (B) and the 6th cycle of LLTC (C). White bar represents light-on, black bar, yellow and blue bars schematise darkness, thermophase and cryophase periods respectively. N(UAS-*cyc*^*DN*^,*Pdf-*GAL80/+): 34, N(*Mai179,Pdf*-GAL80 > *cyc*^*DN*^): 48, N(*Mai179*-GAL4/+): 32. D-E) D-E) Box plot of the slope of the evening peak in the last day of LD (D) and the 6^th^ cycle of LLTC (E). Same flies as in B-C. Statistical test: Kruskal wallis. *: p<0.05, **:p<0.005, ***:p<0.001.

### Morning and evening oscillator neurons are not required for temperature entrainment of the other clock neurons

*Mai179*-gal4 is exclusively expressed in M and E CRY^+^ clock neurons. How the oscillating environmental temperature information entrains the molecular clock in the brain is not known. Previous observations suggest that temperature entrainment uses multiple molecular and probably neuronal circuits involving peripheral thermosensors(Chen et al. 2015; Roessingh et al. 2019; Chen et al. 2018). We therefore tested, whether the *Mai179* cells are capable of entraining the other clock cells in the brain. For this, we dissected brains and quantified PER levels on the 6^th^ day of LLTC at 4 different time points. First, we confirmed the dominant action of *cyc*^*DN*^ in the Mai179 cells and indeed observed only 3/6 LNd at ZT0 (Fig. 8A), while PER levels were constantly low in the other Mai179 cells (sLNv and 5^th^-s-LNv) (Fig. 8B-C). In the remaining LNd CRY^-^, we observed normal PER oscillations albeit with a slight decrease of the amplitude (Fig. 5C). Normal PER cycling was also maintained in the other neurons not expressing *Mai179* (Fig. 8D-E), indicating that they receive independent temperature input. However, we did notice a reduction of the amplitude or a phase shift of PER oscillation in the lLNv (Fig. 8D), which could be due to an influence from the sLNv. Combined with the *Mai179/Pdf-gal80 > cyc*^*DN*^ results (Figure 7), we therefore conclude that synchronized evening activity in LLTC results exclusively from the presence of a functional molecular clock in the CRY^+^ Mai179 E-cells.

**Figure 8:**
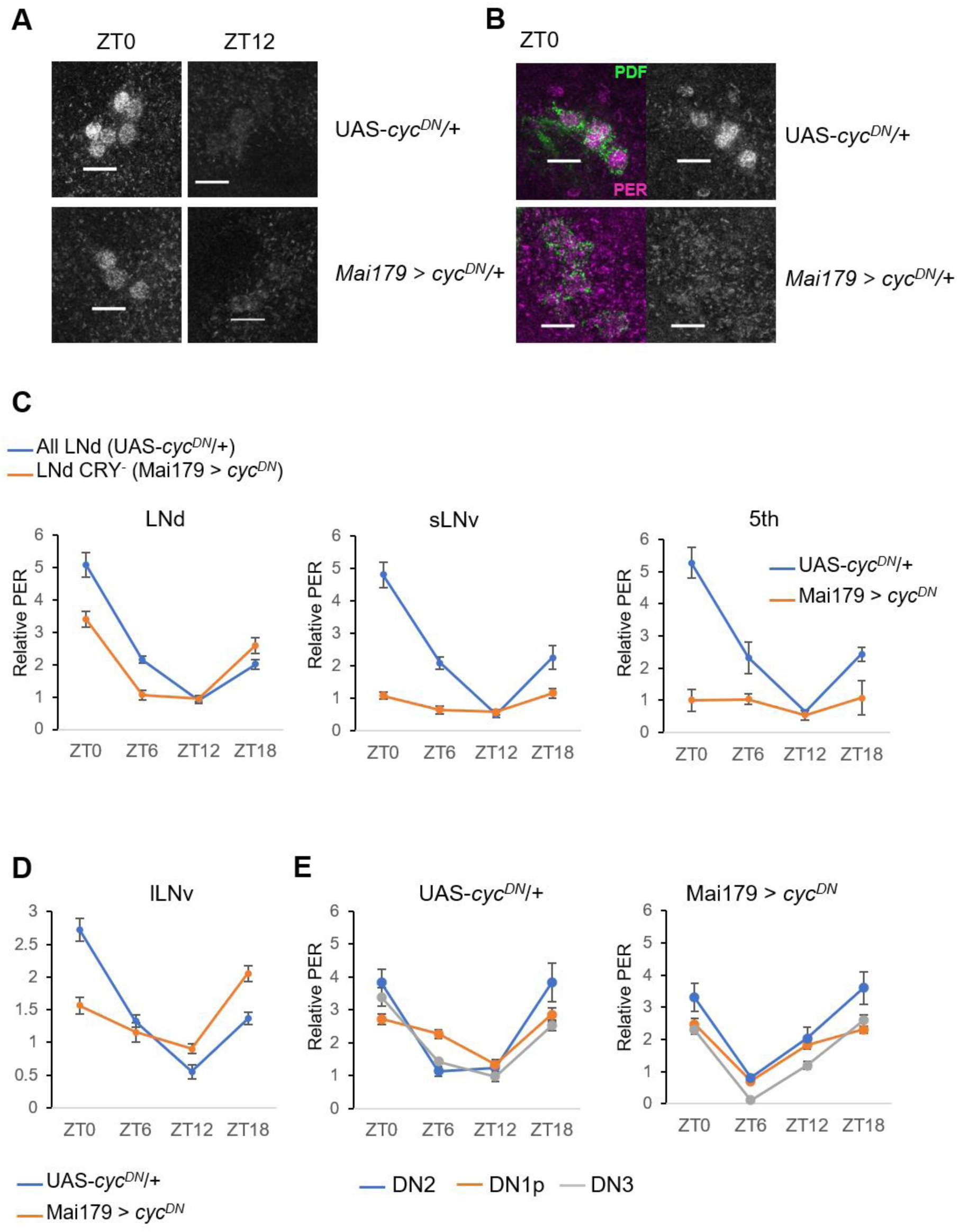
The Mai179^+^ cells do not influence the temperature entrainment of the other clock neurons. A-B) PER immunostaining in LLTC. A) PER in the LNd at ZT0 and ZT12 in controls (UAS-*cyc*^*DN*^/+) and experimentals (*Mai179* > *cyc*^*DN*^). B) PER and PDF immunostaining in the sLNv at ZT0. Left panel: merge images of PDF (green) and PER (magenta). Right panel: PER only. Scale bar: 10µm. C) Average of the relative PER level in Mai179^+^ cells. Since the Mai179^+^ LNd were not found in the experimental flies, only the 3 LNd CRY^-^ cells were quantified in these flies while the 6 LNd were quantified in the controls. D-E) Average of relative PER in Mai179^-^ cells, lLNv (D) and the dorsal neurons (E).

### Neuronal activity of the LNd CRY^+^ evening cells cycles during temperature cycles in constant light

Previous studies reported that in LD the neuronal activity of the LNd cycles with a peak in the middle of the day, even though a discrimination between CRY^+^ and CRY^-^ LNd was not performed(Liang et al. 2016). We have demonstrated the requirement of the LNd CRY^+^ neurons for the synchronised increase of behavioural activity at the end of the thermophase and show that the CRY^-^ LNd are not required (Fig. 3B-C). Therefore, we wanted to measure the neuronal activity of only the CRY^+^ cells during the 6^th^ day of LLTC. For this, we expressed the intracellular Ca^2+^ sensor CaMPARI(Fosque et al. 2015) in the evening cells using *Mai179*-Gal4. We compared ZT0, ZT9 and ZT15 in LLTC. Wild type flies present their highest locomotor activity in LLTC around ZT9, the time where we surprisingly, observed the lowest neuronal activity (Fig. 9). We then measured the neuronal activity of the LNd CRY^+^ expressing *UAS-cyc*^*DN*^. This resulted in disturbed Ca^2+^ oscillations, indicating that the rhythmic neuronal activity of the LNd CRY^+^ depends on the presence of a functional clock in either the LNd CRY^+^ themselves or the other Mai179^+^ cells.

**Figure 9:**
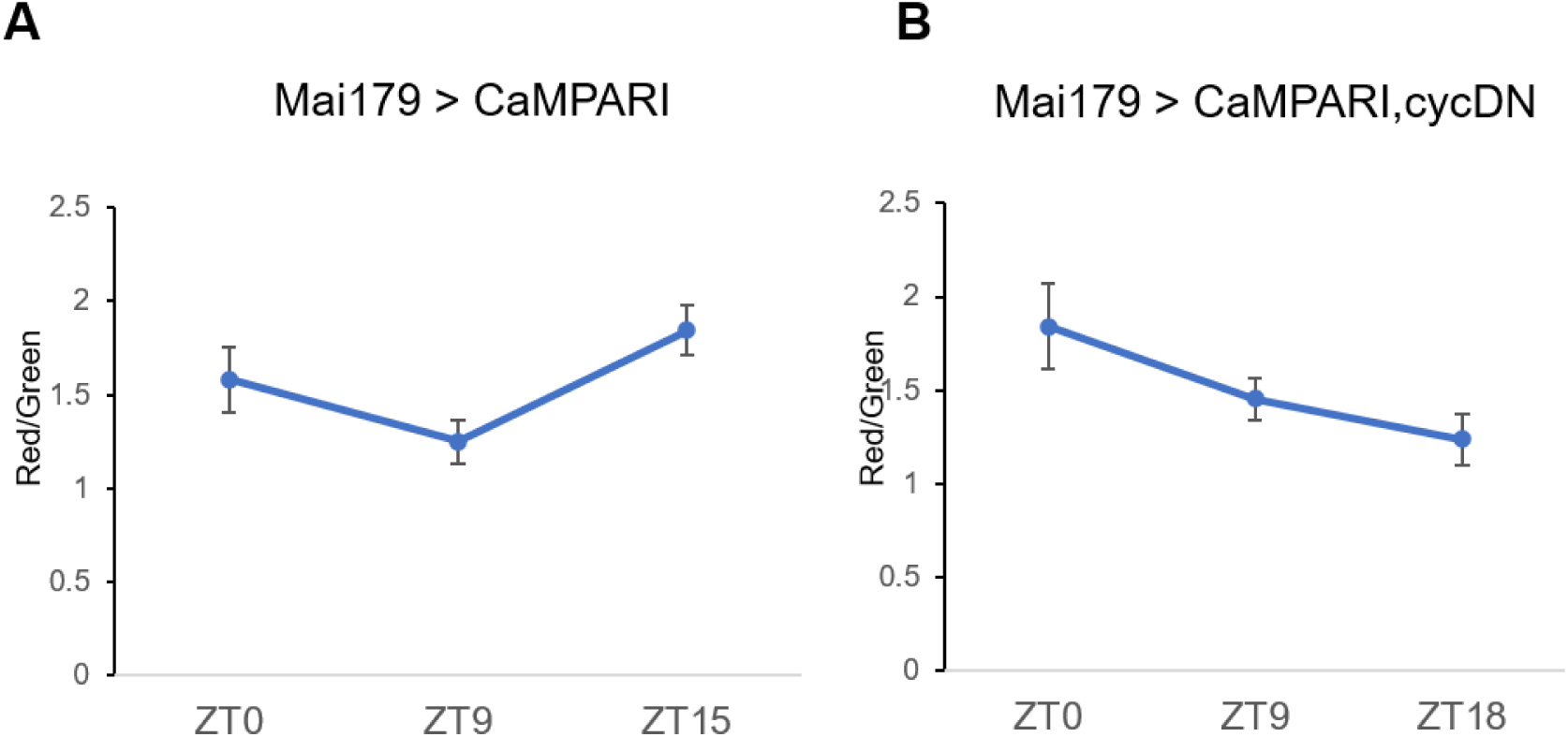
The neuronal activity of the LNd CRY^+^ neurons cycles in LLTC. CaMPARI level of LNd CRY^+^ neurons during the 6^th^ cycle of LLTC in presence (A) and absence (B) of a functional clock in the Mai179^+^ cells.

## Discussion

The daily pattern of locomotor behavior is highly plastic and not only depends on the time of day but also the current environmental condition. How the central nervous system integrates light, temperature and circadian information to drive a locomotor activity is not clear. Here, comparing neuronal function and clock requirement in different environmental conditions, we demonstrated the deterministic role of the E-cells in controlling locomotor output.

The output of the morning oscillator, composed of the PDF^+^ CRY^+^ sLNv, is inhibited by light (Picot et al. 2007), and the morning peak is also repressed by low temperature (Majercak et al. 1999, Zhang et al 2010). Hence, the lack of a morning peak at the end of the cryophase in LLTC 25°C-16°C is therefore not surprising. The evening peak of locomotor activity observed during LLTC occurs almost at the same time as the evening peak in LD. Therefore, the necessity of the E-cells in controlling locomotor activity in LLTC supports the idea that one neuronal oscillator operates under different cyclic environmental conditions. Interestingly, the *acute* environmental parameters at the respective activity peak times are identical between both environmental cycles (light, 25°C), suggesting that the E-cells support behavioral activity under these conditions.

The DN1p clock neurons form a heterogeneous group composed of CRY^+^ and CRY^-^ cells, controlling morning and evening behavioral activity peaks, respectively (Chatterjee et al. 2018), also suggesting a dedicated function for a specific neuronal oscillator. Recently, we have demonstrated that at least a subset of the DN1p directly receive warm temperature input from dTrpA1-expressing cells (Lamaze et al. 2017) and are required for the morning delay of the siesta onset at high temperatures (Lamaze et al. 2017). TrpA1, however, does not play a role in temperature entrainment (Roessingh et al. 2015). Hence, we have to distinguish between neuronal pathways used to synchronize clock neurons from those affecting behavior in response to environmental signals and the time of day. Although the DN1p and their outputs are highly sensitive to temperature changes (Lamaze et al. 2017; Yadlapalli et al. 2018; Zhang, Liu, et al. 2010), as we show here they are not required for controlling the locomotor behavior in LLTC 25°C-16°C. Probably, the reason for this is that the high light intensity used in our study (700lux-1000lux) suppresses the DN1p evening output (Zhang et al. 2010). However, the DN1p are sufficient, along with the other dorsal neurons and the LNd CRY^-^, for driving a rhythmic locomotor behaviour under DDTC 29°C-20°C(Gentile et al. 2013).

Synchronization of clock protein oscillations in LPN neurons is highly sensitive to temperature entrainment (Yoshii et al. 2005; Yoshii et al. 2010; Miyasako et al. 2007). However, we have previously observed that these neurons are not necessary for rhythmic behavior under TC conditions in both LL and constant darkness (Gentile et al. 2013). Here, we confirm their non-requirement for rhythmic locomotor behavior in LLTC, suggesting that LPN sensitivity for molecular synchronization to temperature cycles serves another function, which is not necessarily observable in our laboratory conditions.

CRY is expressed in about half of the clock network in the brain and plays an important role in light entrainment (Helfrich-Förster et al. 2001), consistent with the idea that CRY^+^ neurons are more sensitive to light and CRY^-^ cells are more sensitive to temperature(Yoshii, Hermann, and Helfrich-Förster 2010). However, here we demonstrate an essential role of the CRY^+^ neurons in controlling rhythmic behavior in LLTC. Furthermore, the fact that in the absence of a clock in the Mai179^+^ cells other clock neurons can be synchronized, suggest that multiple and independent pathways contribute to temperature entrainment(Roessingh et al. 2019; Chen et al. 2015; Yadlapalli et al. 2018). In addition to light-resetting, CRY promotes neuronal activity in presence of light (Fogle et al. 2011). Surprisingly however, a functional clock in subsets of the DN and the CRY^-^ LNd neurons is sufficient to drive rhythmic locomotor behavior in LLTC only in the absence of CRY(Gentile et al. 2013). This suggests that the presence of CRY is enough to inhibit the output of the CRY^-^ cells in presence of light.

Clock neurons are highly heterogenous, even inside the same group. However, they can all be entrained by light or by temperature. This heterogeneity suggests that they serve specific functions. Interestingly, although the 5^th^ and 1 LNd CRY^+^ neuron are not necessary for regulating locomotor behavior in LLTC, reducing the activity or stopping the clock in the 2 DN1a and 2 LNd CRY^+^ cells (*R16C05-gal4*: Figures 3A, 6B) is enough to affect rhythmic locomotion. Nonetheless, the effect is weaker compared to a knock-down in all E-cells, suggesting redundancy within this neuronal group.

Knowing that the molecular clock cycles with the same phase in all clock neurons, how can we explain the timing specificity of a locomotor output? According to one model, although the molecular clock is locked to one phase, the neuronal activity cycles with multiple phases(Liang et al. 2016). For example, while the neuronal activity of the sLNv cycles with a peak in the late night/early morning, the LNd cycle with a peak in the late afternoon. These peaks would therefore suggest that both groups are pro-arousal since their phase of activity corresponds to the phase of the locomotor activity they control(Liang et al. 2016). However, this model does not distinguish between CRY^+^ and CRY^-^ LNd. Similarly, the model does not take into account the diversity of functions within the same anatomical group of other clock neurons, for example the DN1p, which indeed, can drive both morning and evening anticipation(Zhang et al. 2010). Recent findings indicate that the CRY^+^ DN1p control mainly the morning peak, while the CRY^-^ DN1p control mainly the evening peak(Chatterjee et al. 2018), questioning if the neuronal activity of all DN1p indeed cycle with the same phase (Liang et al. 2016; Lamaze et al. 2018). Here, we selectively monitored the neuronal activity of the CRY^+^ LNd in LLTC and found a trough of their activity at a time when flies are the most active in LLTC. Recently, another group focused on the DN1a function and physiology (Alpert et al. 2020). They observed that the neuronal activity of these neurons cycles with a higher activity during the day. Furthermore, these neurons are inhibited by cooling under the preferred temperature (25°C), suggesting a higher amplitude of oscillation in LLTC 25°C-16°C (Alpert et al. 2020). Hence, we can propose an alternative model where the evening cells are less active in the afternoon and therefore inhibit locomotion.

The evening peak observed in standard LD condition is controlled by the evening cells, however, the phase of this event is controlled by the lLNv via PDF signaling(Cusumano et al. 2009; Schlichting et al. 2019). Nonetheless, a clock in these cells is not necessary for the evening output to happen. Although we do not know the environmental conditions required to advance the phase of the evening peak in a PDF-dependent manner, the visual system, via the rhodopsins Rh5 and Rh6 play a role in phasing the evening locomotor activity(Schlichting et al. 2019; Cusumano et al. 2009). As discussed above, the acute environmental condition during the thermophase in LLTC, is not different from the one in LD during the light phase, explaining why the LNd evening cells are important in both cases. However, we did notice that the phase of this event is advanced in LLTC compared to LD, mimicking the *Pdf*^*0*^ behavioral phenotype observed under standard LD conditions (Renn et al. 1999) or that of wild type flies in LD and low temperatures (Majercak et al. 1999). This intriguing observation suggests a potential role of the visual system and PDF in phasing the evening output under the presence of light and different night/cryophase temperatures.

Seasonal adaptation corresponds to the occurrence of a clock-controlled behavior at a specific time of day depending on the current environmental condition. Here, by showing that the same group of neurons drive the same locomotor behavior independently of the Zeitgeber used for their entrainment, suggests that a specific oscillator controls a particular output at a dedicated range of time under the same acute environmental condition. The phase of this behavior will depend on the general environmental circumstances. For example, it is advanced in low night temperature (LD18°C vs LD25°C and LLTC 25°C-16°C vs LD25°C). It is therefore conceivable that these neurons also control evening behaviour under natural conditions, ranging from temperate latitudes to those experiencing constant light above the polar circle. In other words, our results show that independent of the Zeitgeber used to synchronize these neurons, they control locomotor activity during the same period of time, truly justifying their classification as ‘evening cells’.

## Acknowledgments

We thank Taishi Yoshii and François Rouyer for providing fly strains. AL received a *Women in Research* (*WiRe*) (Women in Research, fellowship co-funded by the University of Münster and the Deutsche Forschungsgemeinschaft (DFG). AL and RS are supported by the DFG grant STA-421/7-1..

## References

Alpert, M. H., D. D. Frank, E. Kaspi, M. Flourakis, E. E. Zaharieva, R. Allada, A. Para, and M. Gallio. 2020. ‘A Circuit Encoding Absolute Cold Temperature in Drosophila’, Curr Biol.

Bahn, Jae Hoon, Gyunghee Lee, and Jae H Park. 2009. ‘Comparative analysis of Pdf-mediated circadian behaviors between Drosophila melanogaster and D. virilis’, Genetics, 181: 965–75.

Bulthuis, Nicholas, Katrina R Spontak, Benjamin Kleeman, and Daniel J Cavanaugh. 2019. ‘Neuronal Activity in Non-LNv clock cells is required to produce Free-Running rest: activity rhythms in Drosophila’, Journal of biological rhythms, 34: 249–71.

Chatterjee, Abhishek, Angélique Lamaze, Joydeep De, Wilson Mena, Elisabeth Chélot, Béatrice Martin, Paul Hardin, Sebastian Kadener, Patrick Emery, and François Rouyer. 2018. ‘Reconfiguration of a multi-oscillator network by light in the Drosophila circadian clock’, Current Biology, 28: 2007-17. e4.

Chen, Chenghao, Edgar Buhl, Min Xu, Vincent Croset, Johanna S Rees, Kathryn S Lilley, Richard Benton, James JL Hodge, and Ralf Stanewsky. 2015. ‘Drosophila Ionotropic Receptor 25a mediates circadian clock resetting by temperature’, Nature, 527: 516.

Chen, Chenghao, Min Xu, Yuto Anantaprakorn, Mechthild Rosing, and Ralf Stanewsky. 2018. ‘nocte is required for integrating light and temperature inputs in circadian clock neurons of Drosophila’, Current Biology, 28: 1595-605. e3.

Cusumano, Paola, André Klarsfeld, Elisabeth Chélot, Marie Picot, Benjamin Richier, and François Rouyer. 2009. ‘modulated visual inputs and cryptochrome define diurnal behavior in Drosophila’, Nature neuroscience, 12: 1431.

Fogle, Keri J, Kelly G Parson, Nicole A Dahm, and Todd C Holmes. 2011. ‘CRYPTOCHROME is a blue-light sensor that regulates neuronal firing rate’, Science, 331: 1409–13.

Fosque, Benjamin F, Yi Sun, Hod Dana, Chao-Tsung Yang, Tomoko Ohyama, Michael R Tadross, Ronak Patel, Marta Zlatic, Douglas S Kim, and Misha B Ahrens. 2015. ‘Labeling of active neural circuits in vivo with designed calcium integrators’, Science, 347: 755–60.

Gentile, Carla, Hana Sehadova, Alekos Simoni, Chenghao Chen, and Ralf Stanewsky. 2013. ‘Cryptochrome antagonizes synchronization of Drosophila’s circadian clock to temperature cycles’, Current Biology, 23: 185–95.

Grima, Brigitte, Elisabeth Chélot, Ruohan Xia, and François Rouyer. 2004. ‘Morning and evening peaks of activity rely on different clock neurons of the Drosophila brain’, Nature, 431: 869.

Gummadova, Jennet Orazmuradovna, Graham Andrew Coutts, and Nicholas Robert John Glossop. 2009. ‘Analysis of the Drosophila Clock promoter reveals heterogeneity in expression between subgroups of central oscillator cells and identifies a novel enhancer region’, Journal of biological rhythms, 24: 353–67.

Harper, Ross EF, Peter Dayan, Joerg T Albert, and Ralf Stanewsky. 2016. ‘Sensory conflict disrupts activity of the Drosophila circadian network’, Cell reports, 17: 1711–18.

Helfrich-Förster, Charlotte, Christine Winter, Alois Hofbauer, Jeffrey C Hall, and Ralf Stanewsky. 2001. ‘The circadian clock of fruit flies is blind after elimination of all known photoreceptors’, Neuron, 30: 249–61.

Ho, Joses, Tayfun Tumkaya, Sameer Aryal, Hyungwon Choi, and Adam Claridge-Chang. 2019. ‘Moving beyond P values: data analysis with estimation graphics’, Nature Methods, 16: 565–66.

Johard, Helena AD, Taishi Yoishii, Heinrich Dircksen, Paola Cusumano, Francois Rouyer, Charlotte Helfrich-Förster, and Dick R Nässel. 2009. ‘Peptidergic clock neurons in Drosophila: ion transport peptide and short neuropeptide F in subsets of dorsal and ventral lateral neurons’, Journal of Comparative Neurology, 516: 59–73.

Kaneko, Haruna, Lauren M Head, Jinli Ling, Xin Tang, Yilin Liu, Paul E Hardin, Patrick Emery, and Fumika N Hamada. 2012. ‘Circadian rhythm of temperature preference and its neural control in Drosophila’, Current Biology, 22: 1851–57.

Lamaze, Angélique, Patrick Krätschmer, Ko-Fan Chen, Simon Lowe, and James EC Jepson. 2018. ‘A Wake-Promoting circadian output circuit in Drosophila’, Current Biology, 28: 3098-105. e3.

Lamaze, Angelique, Patrick Kratschmer, and James EC Jepson. 2018. ‘A sleep-regulatory circuit integrating circadian, homeostatic and environmental information in Drosophila’, bioRxiv: 250829.

Lamaze, Angelique, Arzu Öztürk-Çolak, Robin Fischer, Nicolai Peschel, Kyunghee Koh, and James EC Jepson. 2017. ‘Regulation of sleep plasticity by a thermo-sensitive circuit in Drosophila’, Scientific reports, 7: 40304.

Levine, Joel D, Pablo Funes, Harold B Dowse, and Jeffrey C Hall. 2002. ‘Advanced analysis of a cryptochrome mutation’s effects on the robustness and phase of molecular cycles in isolated peripheral tissues of Drosophila’, BMC neuroscience, 3: 5.

Liang, Xitong, Timothy E Holy, and Paul H Taghert. 2016. ‘Synchronous Drosophila circadian pacemakers display nonsynchronous Ca2+ rhythms in vivo’, Science, 351: 976–81.

Liu, S., A. Lamaze, Q. Liu, M. Tabuchi, Y. Yang, M. Fowler, R. Bharadwaj, J. Zhang, J. Bedont, S. Blackshaw, T. E. Lloyd, C. Montell, A. Sehgal, K. Koh, and M. N. Wu. 2014. ‘WIDE AWAKE mediates the circadian timing of sleep onset’, Neuron, 82: 151–66.

Majercak, John, David Sidote, Paul E Hardin, and Isaac Edery. 1999. ‘How a circadian clock adapts to seasonal decreases in temperature and day length’, Neuron, 24: 219–30.

Miyasako, Yoko, Yujiro Umezaki, and Kenji Tomioka. 2007. ‘Separate sets of cerebral clock neurons are responsible for light and temperature entrainment of Drosophila circadian locomotor rhythms’, Journal of biological rhythms, 22: 115–26.

Murad, Alejandro, Myai Emery-Le, and Patrick Emery. 2007. ‘A subset of dorsal neurons modulates circadian behavior and light responses in Drosophila’, Neuron, 53: 689–701.

Picot, M., P. Cusumano, A. Klarsfeld, R. Ueda, and F. Rouyer. 2007. ‘Light activates output from evening neurons and inhibits output from morning neurons in the Drosophila circadian clock’, PLoS Biol, 5: e315.

Pittendrigh, Colin S. 1954. ‘On temperature independence in the clock system controlling emergence time in Drosophila’, Proceedings of the National Academy of Sciences of the United States of America, 40: 1018.

Renn, Susan CP, Jae H Park, Michael Rosbash, Jeffrey C Hall, and Paul H Taghert. 1999. ‘A pdf neuropeptide gene mutation and ablation of PDF neurons each cause severe abnormalities of behavioral circadian rhythms in Drosophila’, Cell, 99: 791–802.

Roessingh, Sanne, Mechthild Rosing, Martina Marunova, Maite Ogueta, Rebekah George, Angelique Lamaze, and Ralf Stanewsky. 2019. ‘Temperature synchronization of the Drosophila circadian clock protein PERIOD is controlled by the TRPA channel PYREXIA’, Communications biology, 2: 246.

Roessingh, Sanne, Werner Wolfgang, and Ralf Stanewsky. 2015. ‘Loss of Drosophila melanogaster TRPA1 function affects “siesta” behavior but not synchronization to temperature cycles’, Journal of biological rhythms, 30: 492–505.

Schlichting, Matthias, Patrick Weidner, Madelen Diaz, Pamela Menegazzi, Elena Dalla Benetta, Charlotte Helfrich-Foerster, and Michael Rosbash. 2019. ‘Light-mediated circuit switching in the Drosophila neuronal clock network’, Current Biology, 29: 3266-76. e3.

Schubert, Frank K, Nicolas Hagedorn, Taishi Yoshii, Charlotte Helfrich-Förster, and Dirk Rieger. 2018. ‘Neuroanatomical details of the lateral neurons of Drosophila melanogaster support their functional role in the circadian system’, Journal of Comparative Neurology, 526: 1209–31.

Sekiguchi, Manabu, Kotaro Inoue, Tian Yang, Dong-Gen Luo, and Taishi Yoshii. 2020. ‘A Catalog of GAL4 Drivers for Labeling and Manipulating Circadian Clock Neurons in Drosophila melanogaster’, Journal of biological rhythms, 35: 207–13.

Stanewsky, Ralf, Brigitte Frisch, Christian Brandes, Melanie J Hamblen-Coyle, Michael Rosbash, and Jeffrey C Hall. 1997. ‘Temporal and Spatial Expression Patterns of Transgenes Containing Increasing Amounts of the Drosophila Clock Geneperiod and a lacZ Reporter: Mapping Elements of the PER Protein Involved in Circadian Cycling’, Journal of Neuroscience, 17: 676–96.

Stoleru, Dan, Ying Peng, José Agosto, and Michael Rosbash. 2004. ‘Coupled oscillators control morning and evening locomotor behaviour of Drosophila’, Nature, 431: 862.

Sweeney, Sean T, Kendal Broadie, John Keane, Heiner Niemann, and Cahir J O’Kane. 1995. ‘Targeted expression of tetanus toxin light chain in Drosophila specifically eliminates synaptic transmission and causes behavioral defects’, Neuron, 14: 341–51.

Tanoue, Shintaro, Parthasarathy Krishnan, Balaji Krishnan, Stuart E Dryer, and Paul E Hardin. 2004. ‘Circadian clocks in antennal neurons are necessary and sufficient for olfaction rhythms in Drosophila’, Current Biology, 14: 638–49.

Yadlapalli, Swathi, Chang Jiang, Andrew Bahle, Pramod Reddy, Edgar Meyhofer, and Orie T Shafer. 2018. ‘Circadian clock neurons constantly monitor environmental temperature to set sleep timing’, Nature, 555: 98.

Yoshii, Taishi, Christiane Hermann, and Charlotte Helfrich-Förster. 2010. ‘Cryptochrome-positive and-negative clock neurons in Drosophila entrain differentially to light and temperature’, Journal of biological rhythms, 25: 387–98.

Yoshii, Taishi, Yoshihiro Heshiki, Tadashi Ibuki-Ishibashi, Akira Matsumoto, Teiichi Tanimura, and Kenji Tomioka. 2005. ‘Temperature cycles drive Drosophila circadian oscillation in constant light that otherwise induces behavioural arrhythmicity’, European Journal of Neuroscience, 22: 1176–84.

Yoshii, Taishi, Takeshi Todo, Corinna Wülbeck, Ralf Stanewsky, and Charlotte Helfrich-Förster. 2008. ‘Cryptochrome is present in the compound eyes and a subset of Drosophila’s clock neurons’, Journal of Comparative Neurology, 508: 952–66.

Zhang, Luoying, Brian Y Chung, Bridget C Lear, Valerie L Kilman, Yixiao Liu, Guruswamy Mahesh, Rose-Anne Meissner, Paul E Hardin, and Ravi Allada. 2010. ‘DN1p circadian neurons coordinate acute light and PDF inputs to produce robust daily behavior in Drosophila’, Current Biology, 20: 591–99.

Zhang, Yong, Yixiao Liu, Diana Bilodeau-Wentworth, Paul E Hardin, and Patrick Emery. 2010. ‘Light and temperature control the contribution of specific DN1 neurons to Drosophila circadian behavior’, Current Biology, 20: 600–05.

